# *In vivo* mRNA delivery to virus-specific T cells by light-induced ligand exchange of MHC class I antigen-presenting nanoparticles

**DOI:** 10.1101/2021.10.14.464373

**Authors:** Fang-Yi Su, Qingyang Zhao, Shreyas N. Dahotre, Lena Gamboa, Swapnil Subhash Bawage, Aaron D. Silva Trenkle, Ali Zamat, Hathaichanok Phuengkham, Rafi Ahmed, Philip J. Santangelo, Gabriel A. Kwong

## Abstract

Simultaneous delivery of mRNA to multiple populations of antigen (Ag)-specific CD8^+^ T cells is challenging given the diversity of peptide epitopes and polymorphism of class I major histocompatibility complexes (MHCI). We developed Ag-presenting nanoparticles (APNs) for mRNA delivery using pMHCI molecules that were refolded with photocleavable peptides to allow rapid ligand exchange by UV light and site-specifically conjugated with a lipid tail for post-insertion into preformed mRNA lipid nanoparticles. Across different TCR transgenic mouse models (P14, OT-1, Pmel), UV-exchanged APNs bound and transfected their cognate Ag-specific CD8^+^ T cells equivalent to APNs produced using conventionally refolded pMHCI molecules. In mice infected with PR8 influenza, multiplexed delivery of UV-exchanged APNs against three immunodominant epitopes led to ~50% transfection of a VHH mRNA reporter in cognate Ag-specific CD8+ T cells. Our data shows that UV-mediated peptide exchange can be used to rapidly produce APNs for mRNA delivery to multiple populations of Ag-specific T cells *in vivo*.

**Teaser:** Light-induced rapid production of antigen-presenting nanoparticles for mRNA delivery to multiple virus-specific T cell populations.

## INTRODUCTION

Antigen (Ag)-specific CD8+ T cells express T cell receptors (TCRs) that recognize processed peptide antigens bound to major histocompatibility complex class I (MHCI) molecules expressed on the cell surface. The TCR-pMHCI interaction forms the basis for the exquisite specificity of CD8+ T cell recognition and their cytotoxic activity against target cells bearing cognate pMHCI antigens. This central mechanism has driven increasing interest in delivery approaches that can target and modulate T cells for immunotherapy. Recent studies include delivery of immunomodulatory molecules (e.g., TGFβ inhibitors) using nanoparticles decorated with antibodies against T cell surface markers, including CD3 and PD-1, to enhance effector functions within the tumor microenvironment (*1-4*). Programming endogenous CD3+ or CD8+ T cells with polymer/lipid nanoparticles loaded with nucleic acids [e.g., CD45 siRNA, chimeric antigen receptor (CAR)-encoded DNA] has shown potential to silence target genes in T cells or for *in situ* manufacturing of CAR T cells (*5-9*). To target Ag-specific T cells *in vivo*, strategies include engineered human pMHCI (human leukocyte antigen, HLA)-Fc fusion dimers to expand human papillomavirus (HPV)-specific CD8+ T cells against HPV-associated malignancies (*10*) or track virus-specific CD8+ T cells by immuno-PET imaging (*11*); tumor-targeting antibodies to deliver viral peptides that are cleaved by tumor proteases and then loaded onto MHCI on the tumor cell surface to redirect virus-specific T cells against tumors (*12, 13*); and nanoparticles decorated with pMHC class II molecules to reprogram autoantigen-reactive CD4+ T cells into disease-suppressing regulatory T cells (*14, 15*). These studies highlight the broad applications of *in vivo* delivery to Ag-specific T cells.

Despite considerable interest, however, multiplexed delivery to distinct populations of Ag-specific T cells remains challenging owing to the complexity of the immune response. For example, over 500 SARS-CoV-2 CD8+ T cell epitopes restricted cross 26 HLA class I alleles have been described so far (*16*). Conventionally, pMHCI molecules are expressed by individual refolding reactions to assemble three components – an invariant light chain, a polymorphic heavy chain, and a peptide ligand – into the heterotrimeric structure of endogenous pMHCI molecules (*17*). This serial process precludes production of large pMHCI libraries (*18*) until the development of peptide exchange strategies mediated by UV light (*17, 19*), temperature (*20*), dipeptides (*21*), or chaperone proteins (*22*). With UV light-mediated peptide exchange, the heavy and light chains are refolded with a sacrificial peptide containing a photolabile group, such that upon photocleavage by UV light, the sacrificial peptide dissociates to allow an exchange peptide to bind to the MHCI presentation groove (*17, 19*). For a particular MHC allele, a single batch of UV-sensitive pMHCI molecules can be conventionally refolded and then used to produce 100s of pMHCI molecules carrying different peptides in one step. For example, pMHCI tetramer libraries with >1,000 peptide specificities have been described for the detection of neoAg-specific T cells (*23*).

Here we developed Ag-presenting lipid nanoparticles (APNs) synthesized using UV light-mediated ligand exchange for multiplexed mRNA delivery to influenza-specific CD8+ T cells (**Fig. 1**). We used UV light-mediated ligand exchange to produce a panel of pMHCI molecules from a sacrificial pMHCI precursor that was site-specifically modified with a lipid tail. This allowed post-insertion after peptide exchange to preformed lipid nanoparticles (LNPs) – which was formulated based on a similar MC3-based composition as the first FDA-approved siRNA drug (Onpattro®) (*24*) – encapsulating a model mRNA reporter encoding a camelid VHH antibody (*25*). We found that APNs decorated with conventionally refolded or peptide-exchanged pMHCI molecules targeted and transfected Ag-specific CD8+ T cells in multiple TCR transgenic mouse models in vivo (P14, Pmel-1 and OT-1) regardless of the MHC allotype (H2-D^b^ for P14 and Pmel, H2-K^b^ for OT-1). In a mouse model of recombinant influenza A viral infection (A/Puerto Rico/8/34 H1N1 modified with GP33 antigen, abbreviated as PR8-GP33), intravenously administration of a 3-plex cocktail of peptide-exchanged APNs (NP366/D^b^, PA224/D^b^, GP33/D^b^) resulted in the simultaneous transfection of the top three immunodominant PR8-GP33 specific T cell populations that was significantly more efficient compared to other major cell populations in the spleen and liver. Our data shows that UV light-mediated peptide exchange allows for parallel production of APNs for multiplexed delivery to Ag-specific CD8+ T cells *in vivo*.

**Fig. 1.**
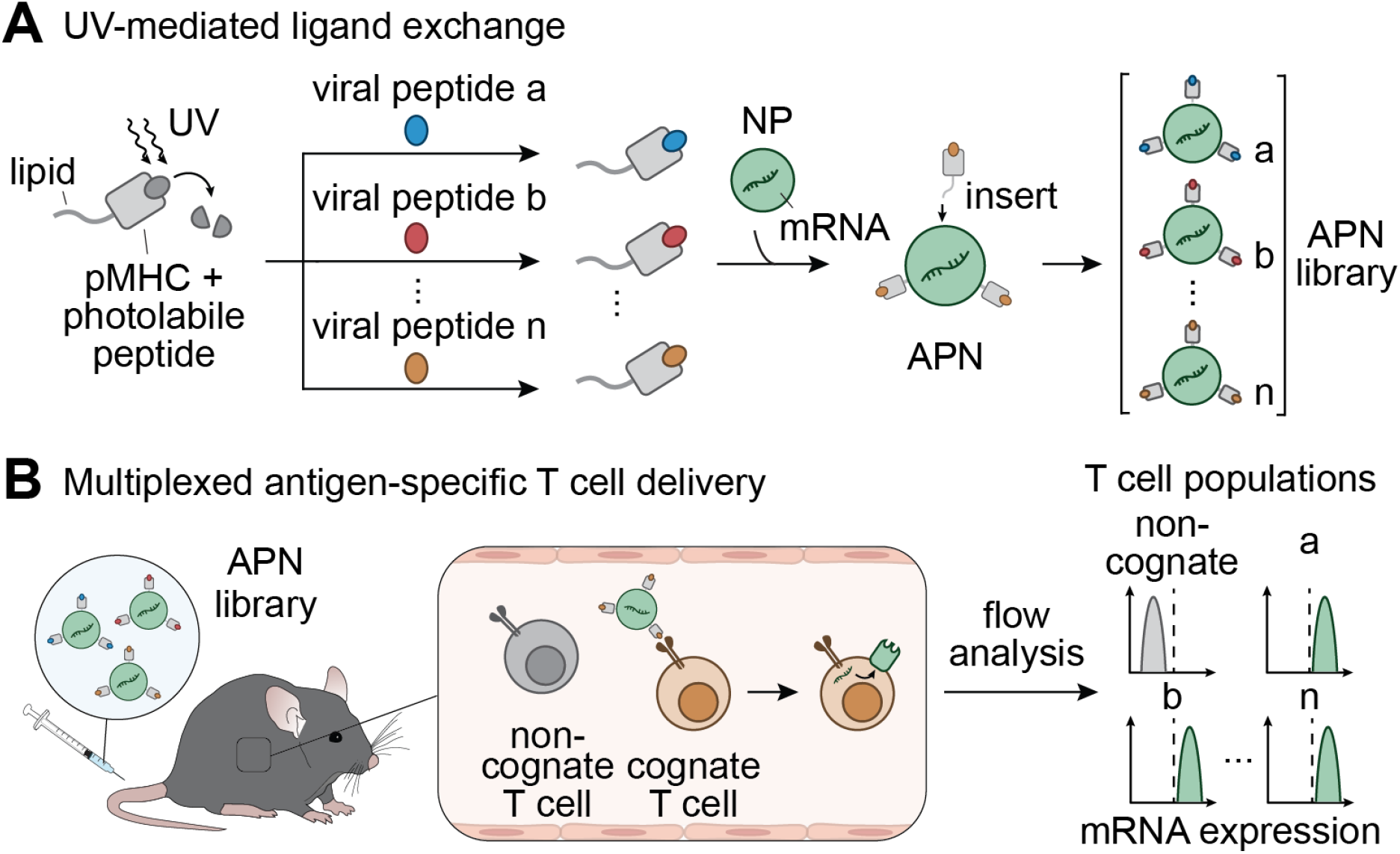
Schematic of UV-mediated peptide exchange of MHC class I antigen-presenting nanoparticles (APNs) for *in vivo* multiplexed delivery to virus-specific T cells. (**A**) We refold peptide major histocompatibility complex class I (pMHC) molecules with photolabile peptides and then conjugated them to a lipid tail to allow subsequent formulation with preformed lipid nanoparticles (NPs). The presence of UV light cleaves the photolabile peptide and induces replacement of the resulting empty MHC groove with a library of viral peptides. After the UV-mediated peptide ligand exchange, we functionalize pMHC molecules on the surface of preformed NPs via post-insertion to form the APN library for multiplexed delivery to virus-specific T cells. (**B**) After intravenous injection into living mice, APNs selectively target cognate T cell populations and transfect them with model mRNA. We validate the mRNA expression using flow analysis.

## RESULTS

### APNs bind to antigen-specific T cells and induce internalization for mRNA transfection

The insertion of derivatives modified with lipids to pre-formed nanoparticles is a well-established approach (*26*) to decorate nanoparticles with ligands that are stabilized by hydrophobic interactions. For example, post-insertion is commonly used to PEGylate liposomes or LNPs using PEG polymers derivatized with lipid tails (*27*). We therefore first sought to express recombinant pMHCI molecules with a site-specific handle for conjugation of a lipid such that the complex could serve as the starting point for peptide exchange prior to post-insertion to LNPs (**Fig. 1A**). We expressed and refolded the lymphocytic choriomeningitis virus (LCMV) antigen GP33/D^b^ (KAVYNFATM/D^b^) with a C-terminal cysteine in the heavy chain to prevent disruption of native disulfide bonds (*28*) and retain proper pMHCI orientation for TCR recognition (*29*). These were then reacted with DSPE-PEG2000-maleimide to generate lipid-modified GP33/D^b^ molecules. In parallel, we synthesized MC3-based LNPs encapsulating eGFP mRNA by microfluidic mixers that were characterized by an average diameter of 93.85 nm and a zeta potential of -30.20 ± 0.5 mV by dynamic light scattering (**Fig. S1A and S1B**). Post-insertion of lipid-modified GP33/D^b^ pMHCI molecules did not appreciably increase LNP size nor alter the zeta potential (107.9 ± 7.34 nm and -22.73 ± 4.7 mV respectively) (**Fig. S1B and S1C**). We also assessed eGFP mRNA concentration using the Ribogreen assay and found that bare LNPs and APNs were comparable (80.20 ± 0.50% to 73.62 ± 0.58%, respectively) (**Fig. S1C**).

We next tested whether GP33/D^b^ APNs can selectively bind to their cognate CD8+ T cells isolated from TCR transgenic P14 mice whose CD8+ T cells express a TCR that specifically recognizes the LCMV GP33/D^b^ antigen (**Fig. 2A**) (*30*). We found that GP33/D^b^ APNs bound to ~ 97% of P14 CD8+ T cells whereas non-cognate GP100/D^b^ (KVPRNQDWL/D^b^) APNs showed minimal staining (3.22%) (**Fig. 2B**). We further tested APN binding using H2-K^b^ restricted OVA/K^b^ (SIINFEKL/K^b^) APNs and observed similar Ag-specific binding when co-incubated with their cognate CD8+ T cells isolated from OT-1 transgenic mice compared to non-cognate NS2/K^b^ (RTFSFQLI/K^b^) APNs (**Fig. 2C**). We next investigated whether the binding of APNs to T cells would induce internalization by T cells given that pMHCI multimers are known to be rapidly taken up by T cells through TCR clustering and receptor-mediated endocytosis at physiological temperatures (*31*). To do this, we studied the fate of APNs after engaging cognate T cells at 37°C compared to 4°C, which is typically used for pMHCI multimer staining to minimize T cell activation and TCR internalization (*32*). We incubated DiD-labeled OVA/K^b^ APNs with OT-1 CD8+ T cells at 4°C and 37°C, followed by an acidic wash to strip un-internalized APNs bound to the TCRs on T cell surface. We found that DiD fluorescence decreased for cells incubated at 4°C, indicating that APNs remained on the cell surface prior to acid wash (**Fig. 2D** and **2E**). For cells treated at 37°C, however, we observed no change in fluorescence, suggesting efficient T cell internalization of APNs. By contrast, we found no binding and internalization of non-cognate GP100/D^b^ APNs by OT-1 CD8+ T cells under all conditions (**Fig. 2E**). To determine whether pMHCI-induced TCR internalization could result in functional mRNA delivery to T cells, we incubated P14 splenocytes with GP33/D^b^ APNs loaded with eGFP mRNA. We observed a dose-dependent eGFP expression by APN-transfected CD8+ T cells [Mean fluorescent intensity (MFI): 397 and 506 for 1 µg and 2 µg mRNA doses, respectively] in contrast to splenocytes treated with PBS (MFI: 153) or free mRNA (MFI: 118) (**Fig. 2F**). Collectively, these results demonstrate that cognate APNs target, bind and induce T cell uptake for functional mRNA delivery *in vitro*.

**Fig. 2.**
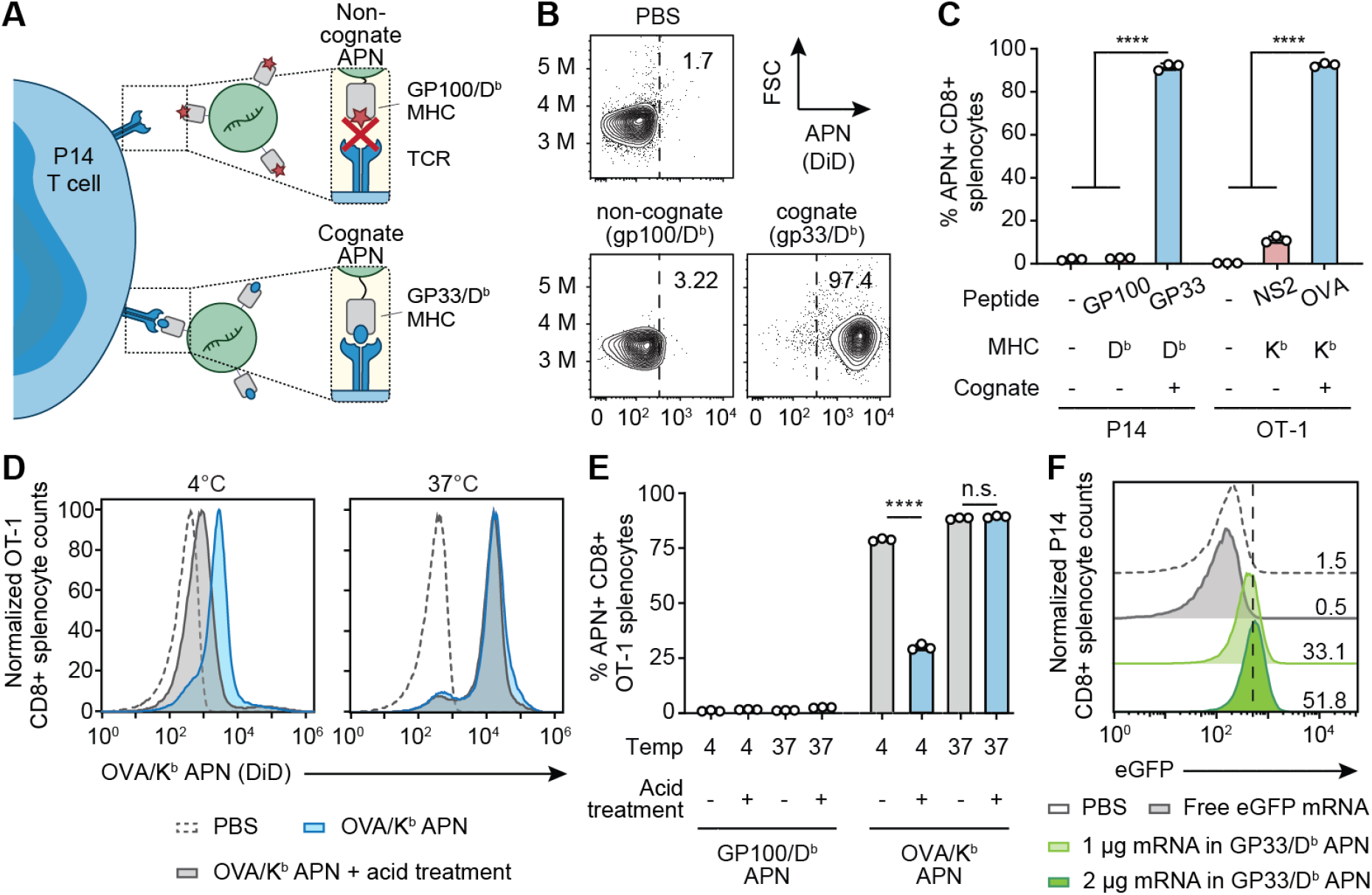
Antigen-presenting nanoparticles (APNs) target antigen-specific T cells and induce cell uptake *in vitro*. (**A**) Illustration of the interaction of P14 CD8+ T cells with its cognate APNs (GP33/D^b^), in contrast to the lack of binding to the non-cognate control (GP100/D^b^ APN). (**B**) Representative flow plots of non-cognate APNs and cognate APNs binding to CD8+ T cells in splenocytes from P14 TCR transgenic mice. Frequencies depicted are based on gating on CD8+ cells. (**C**) Antigen-specific binding of cognate and non-cognate APNs to CD8+ T cells isolated from TCR transgenic P14 or OT-1 mice. * * * * *P* < 0.0001; one-way analysis of variance (ANOVA) and Tukey post-test and correction. All data are means ± SD; n =3 biologically independent wells. (**D**) & (**E**) OT-1 CD8+ splenocytes stained with OVA/K^b^ APNs at 4 or 37 °C and analyzed by flow cytometry before and after treatment with an acetate buffer to strip cell surface proteins. GP100/D^b^ APNs served as a non-cognate control. * * * * *P* < 0.0001 between OVA/K^b^ APN treatment with and without acid treatment at 4°C; n.s. = not significant where *p* = 0.61 between OVA/K^b^ APN with and without acid treatment at 37 °C; two-way ANOVA and Sidak post-test and correction. All data are means ± SD; n =3 biologically independent wells. (**F**) eGFP mRNA expression in P14 CD8+ T cells after co-incubation with free-form eGFP mRNA or eGFP mRNA loaded in GP33/D^b^ APNs for 24 hr.

### APNs transfect antigen-specific T cells in TCR transgenic mice

We next quantified *in vivo* biodistribution and transfection efficiency of GP33/D^b^ APNs in TCR transgenic P14 mice (**Fig. 3A**). We tested functional delivery to major organs using APNs loaded with firefly luciferase (Fluc) mRNA (**Fig. 3B and C**) and found significantly higher luminescence in the spleens isolated from mice treated with GP33/D^b^ APNs compared to non-cognate GP100/D^b^ APNs. Notably, no significant difference was observed in the other major organs between the two groups. To quantify delivery to T cells, we harvested P14 splenocytes 24 hours after infusion of DiD-labeled APNs encapsulating VHH mRNA and observed that cognate GP33/D^b^ APNs targeted >95% of P14 CD8+ T cells while GP100/D^b^ APN controls resulted in <2% binding as quantified by DiD fluorescence (**Fig. 3D**). To quantify functional delivery, we stained for surface expression of VHH and found that GP33/D^b^ APNs resulted in significantly higher transfection efficiency compared to its non-cognate counterpart (~40% versus <2% respectively, **Fig. 3E and F**). Notably, we observed negligible transfection (< 2%) of CD8-splenocytes in mice treated with either GP33/D^b^ APNs or GP100/D^b^ APNs. Combined with the results we observed at the organ level by IVIS imaging (**Fig. 3B and C**), our data suggested that the observed Fluc luminescence in the spleen was mainly from the transfection of cognate CD8+ splenocytes. We further confirmed our results in a different model using Pmel mice that have been engineered to express the cognate TCR against GP100/D^b^. Similar to our results in P14 mice, we observed ~30-40% *in vivo* transfection of Pmel CD8+ splenocytes treated with GP100/D^b^ APNs compared to ~3% transfection with GP33/D^b^ non-cognate APNs (**Fig. S2**). Together, these results demonstrate that APNs enable T cell targeting and functional mRNA delivery in an antigen-specific manner.

**Fig. 3.**
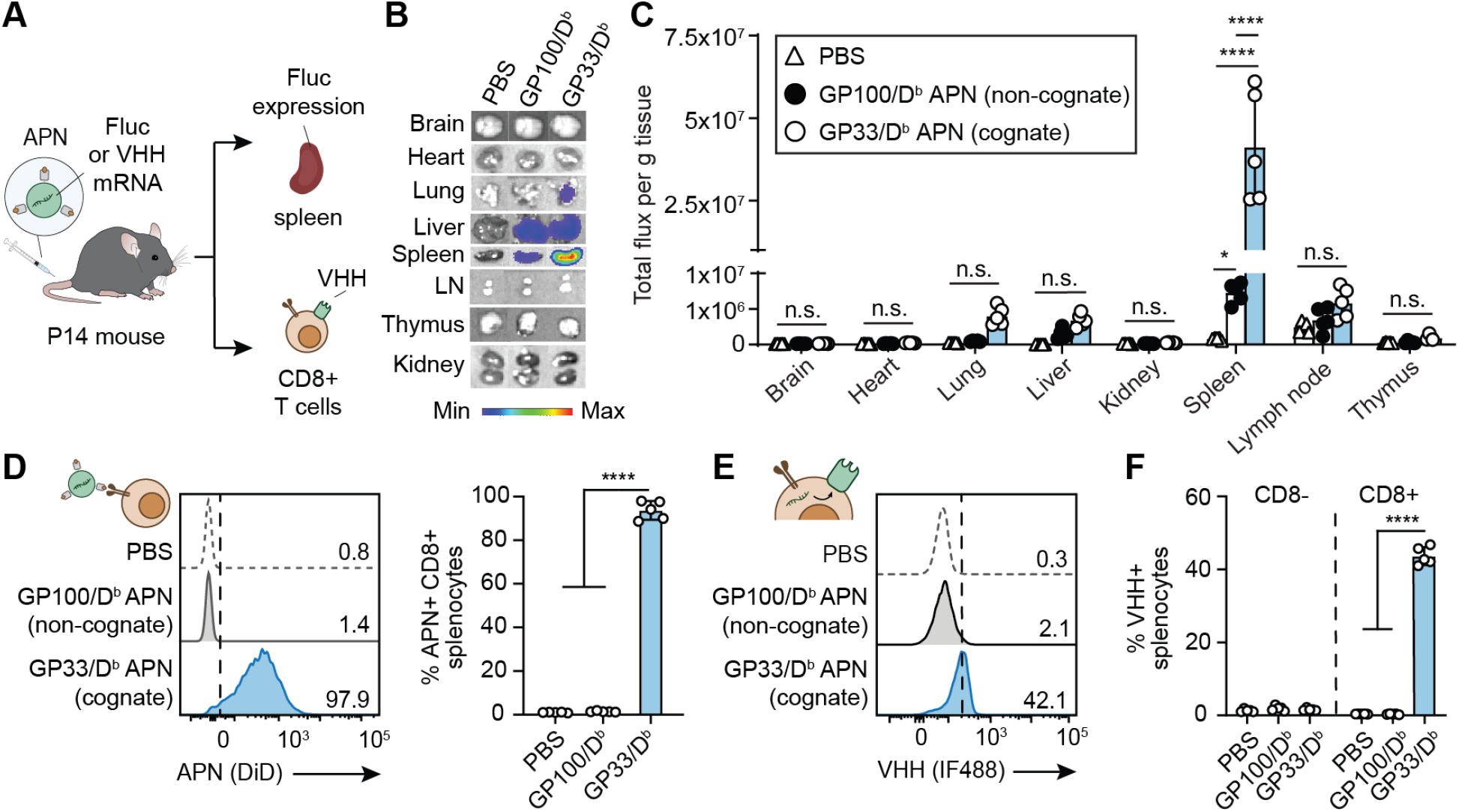
APNs target and transfect antigen-specific T cells in TCR transgenic P14 mice. (**A**) Intravenous injection of GP33/D^b^ APNs to TCR transgenic P14 mice. mRNA encoding firefly luciferase (Fluc) or camelid antibody VHH was loaded in APNs as a reporter. Major organs or splenocytes were harvested for IVIS imaging of Fluc expression or for flow analysis of VHH protein expression on the CD8+ splenocytes. GP100/D^b^ APNs were used as a non-cognate control. (**B**) Representative bioluminescence images of various organs were recorded after 6 h after APN injection to P14 mice. (**C**) Quantification data of bioluminescence images show in (**B**). (**D**) *In vivo* targeting of GP33/D^b^ APNs to CD8+ splenocytes in P14 mice. n.s. = not significant; * *P* = 0.0132 between PBS and GP100/D^b^ APNs; * * * * *P* < 0.0001 between PBS and GP33/D^b^ APNs as well as between GP100/D^b^ APNs and GP33/D^b^ APNs; one-way ANOVA and Tukey post-test and correction. (**E**) Representative flow plot showing *in vivo* APN-mediated transfection in P14 CD8+ splenocytes. (**F**) *In vivo* APN-mediated transfection in both P14 CD8- and CD8+ splenocytes. * * * * *P* < 0.0001 between PBS and GP33/D^b^ APNs as well as between GP100/D^b^ APNs and GP33/D^b^ APNs; one-way ANOVA and Tukey post-test and correction. All data are means ± SD; n = 5 biologically independent mice.

### APNs synthesized by UV-mediated ligand exchange transfect T cells equivalently to folded APNs

Sacrificial peptides that contain a photolabile amino acid and stabilize the MHCI complex during refolding have been previously developed for prevalent alleles including H2-K^b^ and H2-D^b^ in mice (*17, 33*). Therefore, we validated UV-mediated peptide exchange by comparing staining of P14 splenocytes using fluorescent GP33/D^b^ tetramers where the pMHCI monomers were either produced by peptide exchange from ASNENJETM/D^b^ (J represents photocleavable amino acid) or conventionally refolded. We found that both tetramers stained >95% of CD8+ P14 splenocytes, whereas non-cognate GP100/D^b^ tetramers produced by refolding or UV exchange resulted in minimal binding (**Fig. S3**). We further tested UV-exchanged tetramers to detect endogenous immune responses where T cells have a broad range of antigen-specificity and binding affinities to their cognate antigens. To do this, we used the well-characterized mouse model of influenza virus PR8 (A/PR/8/34) modified to express the LCMV GP33 antigen (abbreviated as PR8-GP33). Infection of mice with PR8-GP33 leads to CD8+ T cell responses against at least 16 PR8-derived peptide epitopes including NP366 (ASNENMETM/D^b^), PA224 (SSLENFRAYV/D^b^), PB1-703 (SSYRRPVGI/K^b^), PB1-F2 (LSLRNPILV/D^b^), and NP55 (RLIQNSLTI/D^b^), as well as against GP33 (*34-37*) which served as a positive control epitope. We produced a panel of pMHCI tetramers against these 6 epitopes by peptide exchange (**Fig. 4A and 4B**) (*38*) and validated Ag-specific splenic T cell responses 10 days post-infection (**Fig. 4C**) (*37, 39*).

**Fig. 4.**
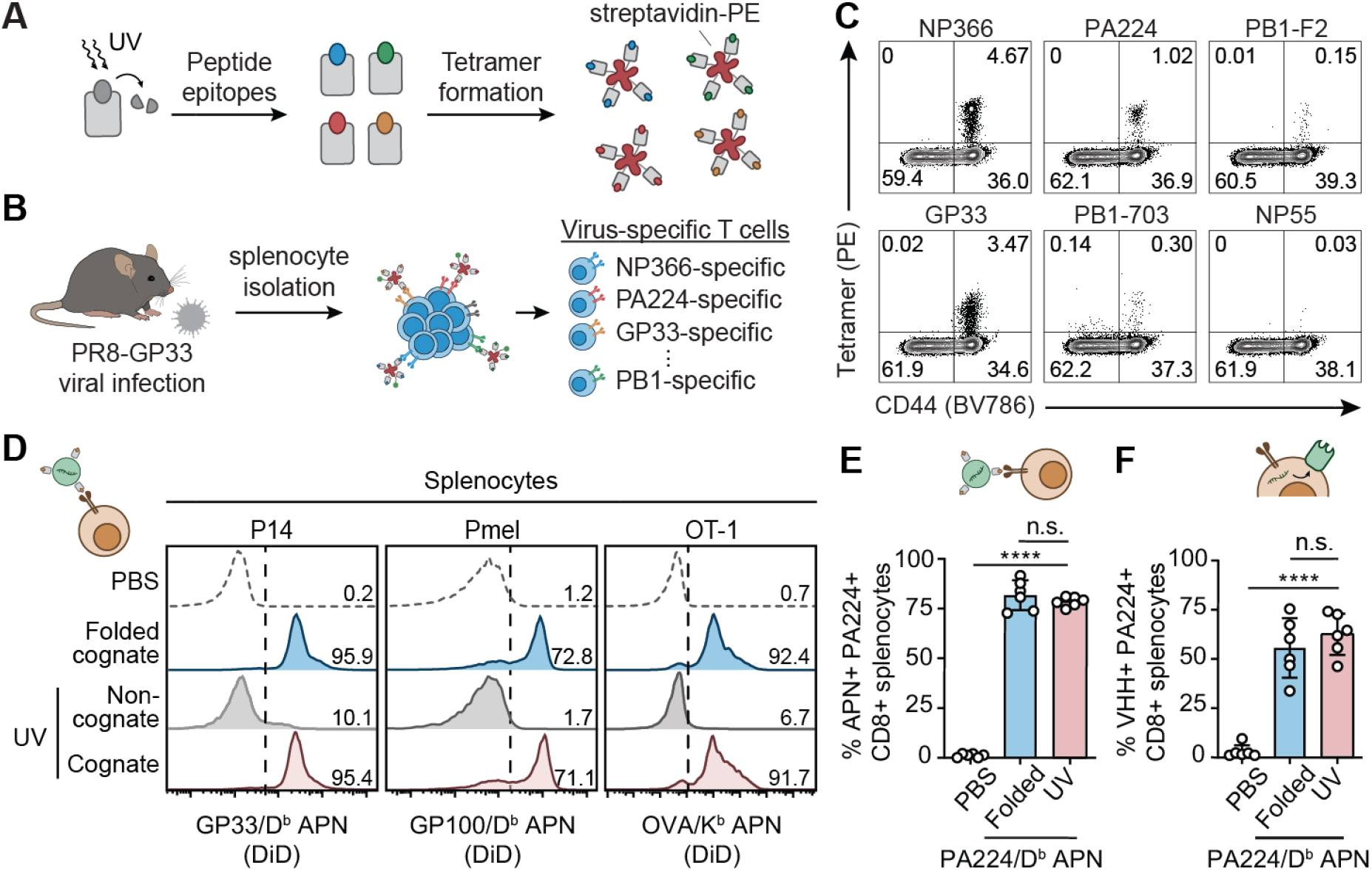
UV-exchanged APNs transfect antigen-specific T cells equivalently to folded APNs. (**A**) Light-triggered peptide exchange technology for high-throughput production of pMHC molecules with various peptide epitopes. pMHC molecules were folded with photolabile peptides that can be cleaved and exchanged with target peptides, followed by tetramer formation with streptavidin conjugated with PE.0 (**B**) Using the UV-exchanged tetramer library to stain virus-specific T cells in a mouse model of PR8-GP33 flu infection. (**C**) Flow cytometry validation of 5 epitopes showing diverse antigen specificity of T cell responses to PR8-GP33 flu infection. (**D**) Equivalent CD8+ splenocyte binding efficiency of UV-exchanged APNs to conventionally folded APNs in three TCR transgenic mouse models *in vitro*. Numbers indicate the percentage of APN+ cells of CD8+ cells. (**E**) *In vivo* targeting activity of UV-exchanged PA224/D^b^ APNs and folded PA224/D^b^ APNs to PA224-specific CD8+ T cells in a mouse model of PR8 infection. n.s. = not significant where *P* = 0.4548 between folded and UV-exchanged PA224/D^b^ APNs; * * * * *P* < 0.0001 between PBS and UV-exchanged PA224/D^b^ APNs; one-way analysis of variance (ANOVA) and Tukey post-test and correction. (**F**) *In vivo* transfection efficiency of UV-exchanged PA224/D^b^ APNs and folded PA224/D^b^ APNs to PA224-specific CD8+ T cells in a mouse model of PR8 infection. n.s. = not significant where *P* = 0.5191 between folded and UV-exchanged PA224/D^b^ APNs; * * * * *P* < 0.0001 between PBS and UV-exchanged PA224/D^b^ APNs; one-way ANOVA and Tukey post-test and correction. For (**E**) and (**F**), all data are means ± SD; n = 6 biologically independent mice.

We next sought to integrate UV-exchange for APN production. To do this, we synthesized a panel of three UV-exchanged APNs inserted with GP33/D^b^, GP100/D^b^, and OVA/K^b^ pMHCI molecules to compare with APNs inserted with pMHCI molecules synthesized using the conventional refolding protocol. In splenocytes isolated from three strains of transgenic mice (P14, Pmel, and OT-1), we found that cognate UV-exchanged APNs bound to CD8+ T cells similar to the folded APNs (**Fig. 4D**). By contrast, we only observed minimal background staining from the non-cognate control APNs. Last, we tested whether the UV-exchanged APNs can target and transfect virus-specific T cells *in vivo* using PR8-infected mice. To do this, we intravenously injected DiD-labeled PA224/D^b^ APNs to PR8-infected mice (*37, 39*). At 24 hr after injection, we found that, consistent with the *in vitro* staining results (**Fig. 4D**), both folded and UV-exchanged PA224/D^b^ APNs targeted ~80% of PA224-specific T cells (**Fig. 4E**) and resulted in comparable transfection efficiency of the model VHH mRNA (**Fig. 4F**).

### Multiplexed T cell transfection using a mouse model of PR8 flu infection

Antibodies against T cell surface markers, including CD3 and CD8, have been used to target polymeric nanoparticles to T cells *in vivo* irrespective of Ag-specificity (*40, 41*). We therefore examined the ability of APNs to transfect virus-specific T cells compared to non-cognate cell populations (**Fig. 5A**). We focused our analysis on liver and spleen as these were the major organs that showed APN accumulation after intravenous administration (**Fig. S4**). Flow cytometry analysis of the major cell types [natural killer cells (NK cells), B cells, CD4 T cells, dendritic cells, macrophages, monocytes, PA224+ flu-specific CD8 T cells, non-cognate PA224-CD8 T cells, Kupffer cells, hepatocytes, and endothelial cells] revealed that APNs preferentially transfected flu virus-specific T cells (PA224+ CD8 T cells, 59.46 ± 11.81%) compared to non-cognate T cells (PA224-CD8 T cells, 2.63 ± 2.17%), which comprise a population diversity of approximately 10^6^ to 10^8^ (**Fig. 5B**) (*42*). As anticipated, transfection was also observed by the reticuloendothelial system – including monocytes and macrophages in the spleen and Kupffer cells in the liver – but at significantly lower transfection efficiency than that of PA224-specific CD8 T cells (* * * * *P* < 0.0001). Compared to cohorts that received PBS, mice that were given folded or UV-exchanged PA224/D^b^ APNs resulted in similar transfection levels across all cell populations studied, supporting their equivalency.

**Fig. 5.**
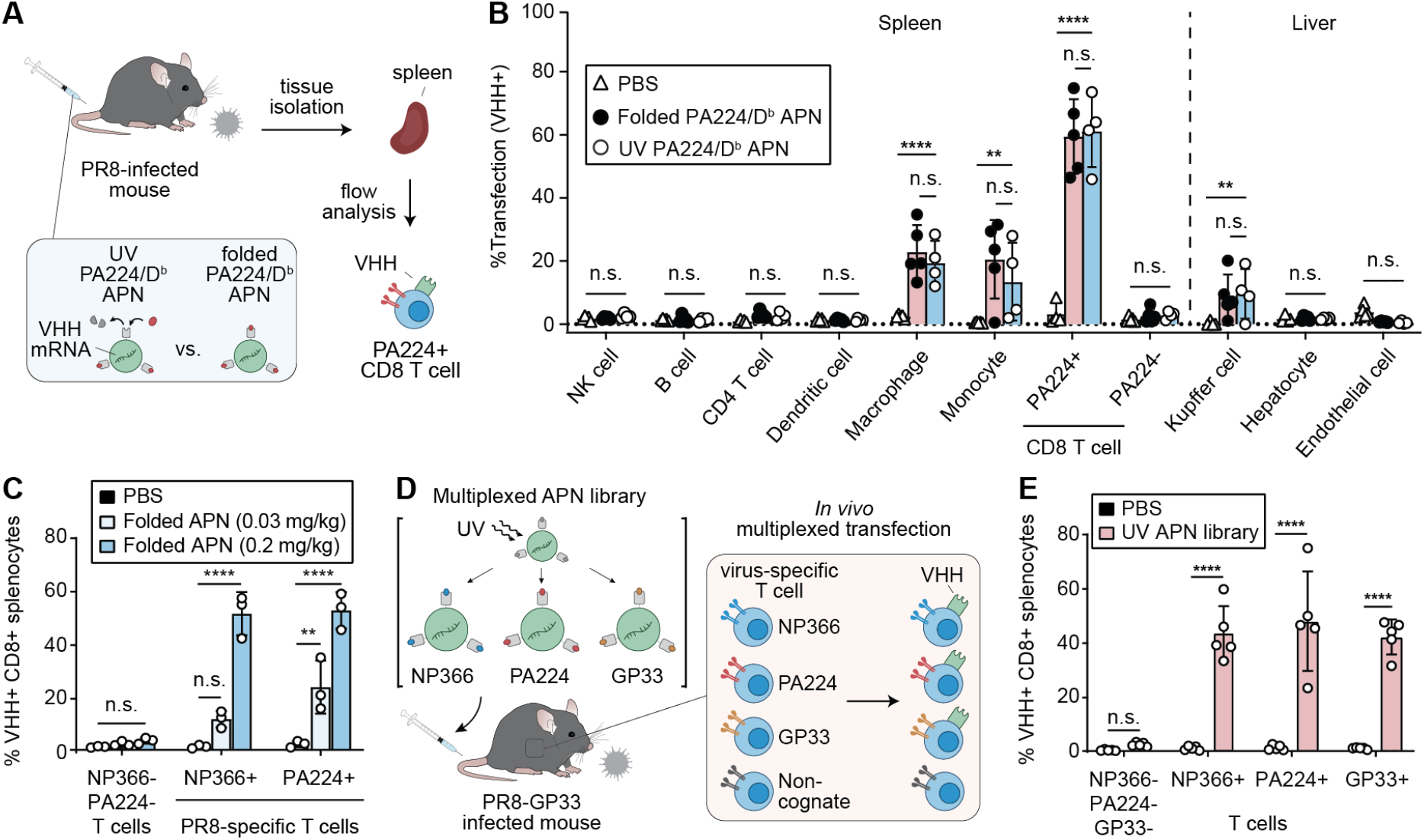
APNs transfect multiplexed T cell subsets with significantly higher transfection efficiency than non-cognate cell populations. (**A**) Schematic of functional biodistribution study at cell level comparing UV-exchanged PA224/D^b^ APNs with folded PA224/D^b^ APNs. (**B**) Transfection efficiency of PBS and PA224/Db APNs in the major cell populations of spleen and liver. n.s. = not significant; * * *P* = 0.0010 between PBS and UV-exchanged PA224/D^b^ APNs in monocyte cell population; * * *P* = 0.0025 between PBS and UV-exchanged PA224/D^b^ APNs in Kupffer cells; * * * * *P* < 0.0001; two-way ANOVA and Dunnett post-test and correction. All data are means ± SD; n = 4-5 biologically independent mice. (**C**) Dose-dependent transfection in two immunodominant flu-specific T cells in PR8 model. Infected mice were treated with a mixture of folded NP366/D^b^ APNs and PA224/D^b^ APNs. n.s. = not significant where *P* = 0.0606 between PBS and folded APNs (0.03 mg/kg total mRNA dose) in NP366+ flu-specific T cell population; * * *P* = 0.0015 between PBS and folded APNs (0.03 mg/kg) in PA224+ flu-specific T cell population (PA224+); * * * * *P* < 0.0001; two-way ANOVA and Sidak post-test and correction. All data are means ± SD; n = 3 biologically independent mice. (**D**) Schematic of multiplexed transfection study. (**E**) Multiplexed transfection study showing the UV-exchanged APN library transfect three virus-specific T cell populations simultaneously. * * * * *P* < 0.0001; two-way ANOVA and Sidak post-test and correction. All data are means ± SD; n = 5 biologically independent mice.

To demonstrate simultaneous transfection of distinct Ag-specific T cells populations *in vivo*, we administered a mixture of DiD-labeled, conventionally refolded NP366/D^b^ and PA224/D^b^ APNs to PR8-infected mice at a mRNA dose of 0.1 and 0.015 mg/kg for each APNs. We found that APNs specifically targeted NP366- and PA224-specific T cells in a dose-dependent manner (**Fig. S5**), whereas no detectable DiD fluorescence was observed in NP366-PA224-T cells (*42*). This resulted in a dose-dependent transfection of NP336+ or PA224+ T cells with the model VHH mRNA (**Fig. 5C**) compared to minimal transfection (<5%) of NP366- and PA224-CD8+ T cells, which was consistent with our earlier observations (**Fig. 5B**). Last, to demonstrate multiplexed transfection with UV-exchanged APNs, we synthesized and pooled a 3-plex panel of APNs presenting the top three immunodominant epitopes (NP366, PA224, GP33) for PR8-GP33 (**Fig. 5D**). This APN library efficiently transfected the three selective clones of T cells with significantly higher expression of the model VHH protein than T cells in mice treated with PBS (**Fig. 5E**). Notably, the transfection efficiency across the three T cell clones were comparable with that of PA224/D^b^ APNs administrated alone (**Fig. 4F and 5B**), suggesting that the APN-mediated multiplexed transfection did not compromise the transfection efficiency in each T cell population tested.

## Discussion

Ag-specific CD8+ T cells are key players in adaptive immunity and their ability to directly kill target cells expressing cognate peptide antigens restricted to MHCI presentation is being harnessed for important applications in cell therapy, vaccines, and autoimmunity (*10-15*). Whereas previous work on delivery to T cells via antibodies against cell surface markers (CD3, CD8, etc.) show great promise, these markers are expressed by all T cells. Moreover, Ag-specific T cell responses are polyclonal (*43, 44*); for instance, across five prevalent HLA-A alleles (HLA-A* 01:01, HLA-A* 02:01, HLA-A* 03:01, HLA-A* 11:01, and HLA-A* 24:02) over 110 flu-specific peptide epitopes have been identified for human influenza A virus (PR8) (*45*). Therefore, we developed APNs for multiplexed mRNA delivery to Ag-specific T cells using UV-mediated peptide exchange to expedite production of APNs against a panel of peptide epitopes. Our *in vivo* data using PR8-infected mice showed that APNs with UV-exchanged pMHCI molecules transfected PA224-specific T cells equivalently to APNs synthesized with conventionally folded pMHCI molecules. This allowed us to construct a 3-plex APN library using UV-mediated peptide exchange to simultaneously transfect the top three immunodominant T cell populations (NP366, PA224, GP33-specific) in a mouse model of PR8-GP33 flu infection.

The use of APNs has the potential to be expanded for more than three peptide epitopes and beyond the two MHC alleles (H-D^b^, H-K^b^) demonstrated in this study. Based on prior studies using the UV-exchange technology to generate pMHCI libraries with thousands of peptide epitopes, we expect that UV-exchange would be sufficient to produce an APN library with 10-20 viral peptide epitopes per MHC allele, the scale we anticipate in common viral infection settings (e.g., CMV, EBV, Flu) (*45-47*). Moreover, sacrificial UV-labile peptides have been developed for most prevalent HLA alleles, including HLA-A* 01:01 (STAPGJLEY), HLA-A* 02:01 (KILGFVFJV), and HLA-A* 11:01 (RVFAJSFIK) (*17, 48, 49*). Therefore, APNs are amenable to other pMHCI molecules, including HLA expressed by human CD8^+^ T cells. The capability of APNs in transfecting multiple virus-specific T cell populations may be used to induce *in vivo* proliferation of virus-specific T cells to treat virus-mediated cancers. For instance, a fusion protein composed of dimerized pMHCI and IL-2 has been developed to expand HPV16 E7_11-20_-specific CD8+ T cells to treat HPV-mediated cancers (*10*), and a recent study suggests that HPV-specific T cells recognizing peptide epitopes derived from HPV E2 and E5 proteins should also be considered to elicit maximal tumor-reactive CD8+ T cell responses against HPV-positive head and neck cancer (*50*).

Other than CD8+ T cells, it may be possible to expand the transfection capability of APNs to CD4^+^ T cells by generating APNs with pMHC class II (pMHCII). For example, Ag-specific CD4^+^ Treg cells are capable of suppressing pathogenic autoimmunity, and therefore prior studies have developed pMHCII-functionalized nanoparticles to expand Ag-specific Tregs through pMHC-TCR interaction for the treatment of autoimmune diseases (*14, 15*). However, this is likely more challenging as binding of class II-bound peptides is less stable compared to that of class I-bound peptides (*51*), and the interaction between the pMHCII and CD4^+^ T cells is weaker than that between the class I counterpart and CD8+ T cells (*52*). Taken together, our data support that the use of APNs for multiplexed mRNA delivery to virus-specific T cells, which can potentially be expanded to transfect broader antigen-specific T cell subsets.

## METHODS

### Animals

6 to 8-week old female mice were used at the outsets of all experiments. P14 (B6;D2-Tg(TcrLCMV)327Sdz/JDvsJ), Pmel (B6.Cg-Thy1a/Cy Tg(TcraTcrb)8Rest/J), and OT-1 (C57BL/6-Tg(TcraTcrb)1100Mjb/J) transgenic mice were bred in house using breeding pairs purchased from Jackson Lab. C57BL/6 for PR8 viral infections were purchased from Jackson Lab. All animal procedures were approved by Georgia Tech IACUC (Protocol numbers: Kwong-A100191, Kwong-A100193, Santangelo-A100169D).

### pMHCI refolding and purification

Peptides used for pMHC refolding were synthesized in housed using the Liberty Blue Peptide Synthesizer (CEM) and validated using LC-MS (Agilent). To generate pMHC molecules for bioconjugation, codon-optimized gBlocks for H2-D^b^ and H2-K^b^ β2m were purchased from IDT and cloned into pET3a vectors (Novagen). H2-Db and H2-Kb genes were engineered with a C-terminal cysteine by site-directed mutagenesis (NEB), and pMHC molecules were expressed and refolded as described previously (*19*).

### ANP preparation and characterization

Lipids, including DSPC, Cholesterol, DMG-PEG, DSPE-PEG (18:0 PEG2000 PE), and DSPE-PEG (2000)-Mal, were purchased from Avanti Polar Lipids. Ionizable lipid D-Lin-MC3-DMA was purchased from MedKoo Biosciences, Inc.. Fluorescent, lipophilic carbocyanine dye DiD was purchase from ThermoFisher scientific. LNP was synthesized as described previously (*53*). Briefly, lipid mixture containing MC3, DSPC, Cholesterol, DMG-PEG, DSPE-PEG (50:10:38:1.5:0.5 molar ratio) and DiD (1% molar ratio of lipid mix) in ethanol was combined with three volumes of mRNA in acetate buffer (10 mM, pH 4.2, 16:1 w/w lipid to mRNA) and injected into micro fluidic mixing device Nanoassemblr (Precision Nanosystems) at a total flow rate of 12 ml/min (3:1 flow rate ratio aqueous buffer to ethanol). mRNA encoding eGFP or membrane-anchored VHH antibody were kind gifts of Dr. Philip Santangelo. Using mRNA encoding membrane-anchored VHH as a reporter gene allows surface expression of VHH to be detected by immunofluorescence staining with anti-VHH antibodies (*25*). The resultant LNPs were diluted 40X in PBS and concentrated down using Amicon spin filter (10kDa, Millipore).

To functionalize the synthesized LNPs with pMHC, pMHC was first coupled with DSPE-PEG-maleimide and decorated on LNPs via post-insertion (*54, 55*). Briefly, a lipid solution of DSPE-PEG and DSPE-PEG (2000)-maleimide at 4:1 molar ratio was dried under nitrogen and placed in vacuum chamber for 2 h to form a thin film. Lipids were rehydrated in PBS at 6.4 mg/ml in a 60°C water bath for 15 min and sonicated in an ultrasonic bath (Branson) for 5 min. Refolded pMHCI monomers with C-terminal Cysteine were reduced with TCEP (1:3 pMHC to TCEP molar ratio) at 37°C for 2 h and mixed with the lipid mixture at RT overnight at 2:1 pMHC/maleimide molar ratio (*56*). Lipid-modified pMHCI molecules were incubated with LNPs at 1:50 maleimide/D-Lin-MC3-DMA molar ratio at RT for 6 h to incorporate pMHCI onto LNPs. The resultant post-insertion mixture was placed in 1 MDa Float-A-Lyzer (Spectrum) and dialyzed against PBS for 16 h.

The sizes of APNs in PBS were measured by dynamic light scattering with Malvern nano-ZS Zetasizer (Malvern). Final lipid concentration was quantified using a phospholipid assay kit (Sigma). The concentration of conjugated pMHCI was determined by BCA assay kit (Sigma). The mRNA encapsulation efficiency was quantified by Quant-iT RiboGreen RNA assay (Life Technology) as previously described (*7*). Briefly, 50 µl of diluted APNs was incubated with 50 µl of 2% Triton X-100 (Sigma-Aldrich) in TE buffer (10 mM Tris-HCl, 20 mM EDTA) in a 96-well fluorescent plate (Costar, Corning) for 10 mins at 37 °C to permeabilize the particle. Then, 100 µl of 1% Ribogreen reagent in TE buffer was added into each well, and the fluorescence (excitation wavelength 485nm, emission wavelength 528 nm) was measured by plate reader (Biotek).

### Primary T cell isolation and activation

Spleens isolated from P14, Pmel, or OT-1 TCR transgenic mice were dissociated in complete RPMI media [RPMI 1640 (Gibco)+10% fetal bovine serum (FBS; Gibco)+1% penicillin/streptomycin (Gibco)] and red blood cells were lysed using RBC lysis buffer (BioLegend). CD8+ T cells were isolated using a CD8a+ Tcell isolation kit (Miltenyi Biotec). For T cell activation, isolated CD8 T cells were cultured in T cell media [complete RPMI media supplemented with 1X nonessential amino acids (Gibco) + 1 × 10^−3^ M sodium pyruvate (Gibco) + 0.05 × 10^−3^ M 2-mercaptoethanol (Sigma)] supplemented with soluble anti-mouse CD28 (5 µg/ml, BD Pharmingen) and 30 U/ml rhIL-2 (Roche) at 1 × 10^6^ cells/ml in wells coated with anti-mouse CD3e (3 µg/ml, BD Pharmingen).

### *In vitro* T cell binding and transfection by APNs

P14, Pmel and OT-1 CD8+ T cells (1×10^6^ cells per sample) were isolated and incubated with APNs (10 µg/ml) in FACS buffer (1× DPBS + 2% FBS + 1 mM EDTA + 25 mM HEPES) for 30 min at 37°C. Cells were washed three times with 1 mL FACS buffer before analysis on a BD Accuri C6. For validation of *in vitro* transfection, P14 CD8+ T cells were activated for 24 h as described above and resuspended in T cell media + 30 U/ml rhIL-2 (Roche) at 2 × 10^6^ cell/ml. 5 × 10^5^ of cells were co-incubated with GP33/D^b^ APN containing eGFP mRNA (1 µg) in 24 well plates at 37°C. After 4 h, 700 µl of T cell media + 30 U/ml rhIL-2 (Roche) was added to each well. After an additional 48 h incubation, cells were washed three times and stained against αCD8 mAb (clone 53-6.7, BioLegend) at 4°C for 30 min. Cells underwent another two washes with FACS buffer before analysis on BD Accuri C6.

### Quantification of APNs internalization by acid wash

OT-1 CD8+ T cells were isolated as described above and incubated with OVA/K^b^, or GP33/D^b^ APNs at 10 µg/ml and αCD8 mAb (clone 53-6.7, BioLegend) at 4 or 37°C for 30 min. Cells were washed with FACS buffer, and a portion of stained cells was analyzed on a BD Accuri C6. The remaining cells were incubated in an acid wash solution (0.5 M NaCl + 0.5 M acetic acid, pH 2.5) for 5 min to strip cell surface proteins as described previously before reanalysis on a BD Accuri C6 (*56*).

### *In vivo* transfection at organ level using P14 TCR transgenic mice

P14 TCR transgenic mice were injected intravenously with GP33/D^b^ or GP100/D^b^ APNs loaded with mRNA encoding firefly luciferase (0.1 mg/kg). Organs were harvested 6 h after injection and incubated in PBS on ice prior to IVIS analysis. Organs were soaked in D-luciferin solution (2 mM luciferin) in PBS for 5 min. After 5-min incubation, bioluminescence images were collected with a Xenogen IVIS Spectrum Imaging System (Xenogen, Alameda, CA). Same type of organs was separated from other organs and imaged together (i.e., spleens from all treatment groups were imaged together).

### *In vivo* T cell targeting and transfection in TCR transgenic mice

P14 or Pmel TCR transgenic mice were injected intravenously with GP33/D^b^ or GP100/D^b^ APNs loaded with GPI-anchored Camelid VHH antibody mRNA (0.2 mg/kg). Splenocytes were harvested 24 h after injection and stained against αCD8 mAb (clone 53-6.7, BioLegend) and anti-Camelid VHH antibody (Genescript) on ice for 30 min. Cells were washed three times with 1 mL FACS buffer before analysis on a BD Accuri C6. Mice splenocytes were harvested 24 h later and stained against αCD8 mAb (clone 53-6.7, BioLegend), anti-Camelid VHH antibody (clone 96A3F5, Genescript), and pMHC tetramers (streptavidin 2 µg/ml) on ice for 30 min. Epitope pMHC tetramers for staining were synthesized in house by mixing biotinylated pMHC with fluorescently labelled streptavidin at a 4:1 molar ratio (*19*). Cells were washed three times with FACS buffer before analysis on BD Accuri C6. All flow data in this study were analyzed with FlowJo v.10 (TreeStar).

### *In vivo* T cell targeting and transfection in PR8-infected mice

PR8 virus was a kind gift of Dr. Philip Santangelo. PR8-GP33 was a kind gift from Dr. Rafi Ahmed (Emory University) and Dr. E. John Wherry (University of Pennsylvania). 6-8 weeks old PR8 infected C57BL/6 mice were intranasally infected with either PR8 virus or PR8-GP33 recombinant virus, as specified in the result section and figure captions. PR8-infected mice were injected intravenously with NP366/D^b^ and PA224/D^b^ APNs ABNs containing the GPI-anchored Camelid VHH antibody mRNA (0.03 or 0.2 mg/kg) on day 10 after viral infection. 24 h after the injection, splenocytes were harvested as described above for immunofluorescent staining. Cells were stained against tetramers (NP366/D^b^, PA224/D^b^, 0.2 µg streptavidin/staining sample), αCD8a mAb (clone 53-6.7, BD), αNK1.1 mAb (clone PK136, TONBO), αB220 mAb (clone RA3-6B2, TONBO), αCD4 mAb (clone RM4-2, Biolegend), anti-Camelid VHH antibody (clone 96A3F5, Genescript) on ice for 30 min (*57-59*). Antibodies were all used at 1:100 dilutions. Cells were then fixed with IC fixation buffer (Thermo) for the flow analysis (Fortessa, BD).

### Functional distribution of APNs at cellular level

6-8 weeks old PR8 infected C57BL/6 mice were injected intravenously with conventional or UV exchanged PA224/D^b^ ABNs containing the GPI-anchored Camelid VHH antibody mRNA (0.1 mg /kg) on day 10 after viral infection. 24 h after the injection, cells from spleen and liver were harvested as described above. Cells were stained against αCD8a mAb (clone 53-6.7, BD), αNK1.1 mAb (clone PK136, TONBO), αB220 mAb (clone RA3-6B2, TONBO), αCD31 mAb (clone PK136, TONBO), αCD45 mAb (clone 30-F11, Biolegend), αCD4 mAb (clone RM4-2, Biolegend), αCD11b mAb (clone M1/70, Biolegend), αCD11c mAb (clone N418, Biolegend), αLy6c mAb (clone HK1.4, Biolegend), αF4/80 mAb (clone BM8, Biolegend), anti-Camelid VHH antibody (clone 96A3F5, Genescript) on ice for 30 min. Antibodies were all used at 1:100 dilutions. Cells were then fixed with IC fixation buffer (Thermo) for the flow analysis (Fortessa, BD).

### Statistical analysis

Significant differences between control and treatment groups were determined by various statistical analyses. Student’s t test was used for two groups comparison. One-way analysis of variance (ANOVA) was used for multiple groups comparison. Two-way ANOVA was used when there were subgroups in each group. Data represent means ± SD in each figure and table as indicated. Statistical analyses were performed using GraphPad Prism 8.0.2 software (GraphPad Software). * P < 0.05, * * P < 0.005, * * * P < 0.0005, and * * * * P < 0.0001.

## Acknowledgement

This work was funded by the Defense Advanced Research Projects Agency (HR00111920008), NIH Director’s New Innovator Award (DP2HD091793), NIH R01 Research Project Grant (R01CA237210-01), the Shurl and Kay Curci Foundation, and the NSF (ECCS-1542174). F.Y.S. was supported by a postdoctoral fellowship from the Department of Biomedical Engineering at Georgia Tech and the College of Engineering at Peking University, China. L.G. was supported by the Alfred P. Sloan Foundation, the NIH GT BioMAT Training Grant (5T32EB006343) and the NSF GRFP (DGE-1451512). S.N.D. was supported by the NSF Graduate Research Fellowships Program (DGE-1650044) and the NSF Integrative Graduate Education and Research Traineeship (DGE-0965945). A.Z. was supported by the Georgia Tech President’s Fellowship. The authors thank Dr. Yun Min Chang for insightful discussion about the experimental designs. The authors thank Hannah Peck (Santangelo lab) for making mRNA used in this work. This work was performed in part at the Georgia Tech Institute for Electronics and Nanotechnology, a member of the National Nanotechnology Coordinated Infrastructure (NNCI), which was supported by the National Science Foundation (ECCS-1542174). G.A.K. holds a Career Award at the Scientific Interface from the Burroughs Wellcome Fund. This content is solely the responsibility of the authors and does not necessarily represent the official views of the NIH.

## Author contributions

F.Y.S. and G.A.K. conceived of the idea. F.Y.S., L.G.., S.N.D., and G.A.K. designed the experiments. F.Y.S., Q.Z., L.G., S.N.D., S.B., A.Z., P.S., G.A.K. interpreted the results. F.Y.S., Q.Z., L.G., S.N.D., S.B., A.Z., and A.D.S.T. performed the experiments. F.Y.S., Q.Z., and G.A.K. wrote the manuscript.

## Data and materials availability

Data generated or analyzed during this study and all unique materials can be made available by the corresponding author on reasonable request.

## Supplementary Materials

**Supplement figure 1.**
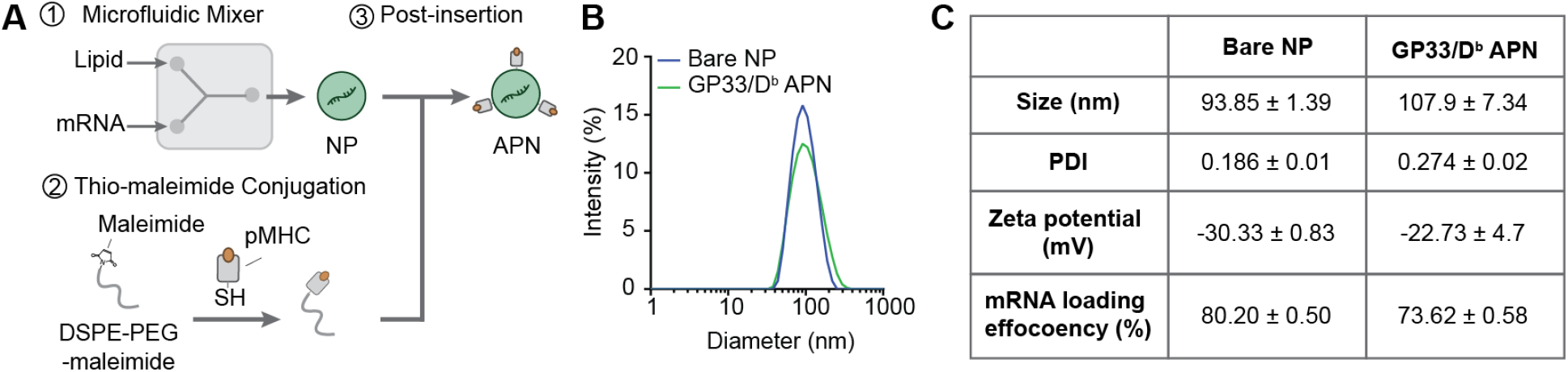
Synthesis and characterization of antigen-presenting nanoparticles (APNs). (**A**) Schematic of the generation of APNs using microfluidic mixers and post-insertion technique. (**B**) Uniform distribution of the synthesized bare nanoparticles (NPs) and APNs functionalized with GP33/D^b^ APN. (**C**) Hydrodynamic size, polydispersity index (PDI), zeta potential, and mRNA loading of the produced NPs. Data are presented as mean ± S.D. n = 3 technical replicates.

**Supplement figure 2.**
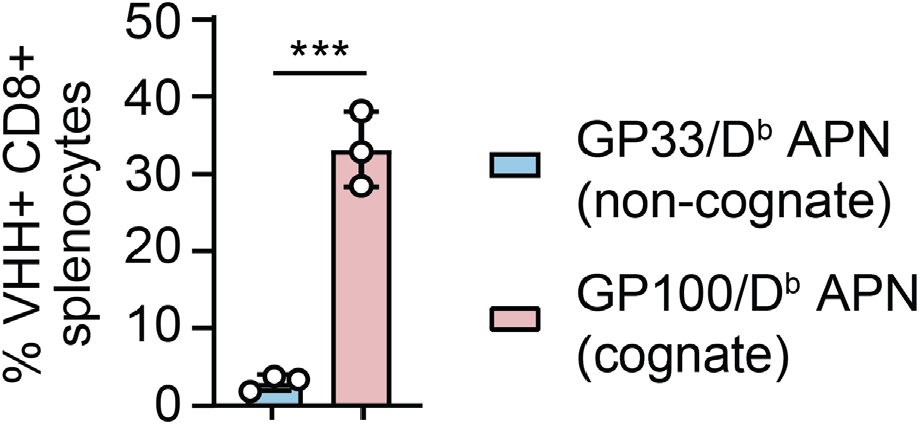
*In vivo* APN-mediated transfection in CD8+ splenocytes isolated from Pmel TCR transgenic mice. * * * *P* < 0.001 between GP33/D^b^ APNs (non-cognate) and GP100/D^b^ APNs (cognate); one-way ANOVA and Tukey post-test and correction. All data are means ± SD; n = 3 biologically independent mice.

**Supplementary figure 3.**
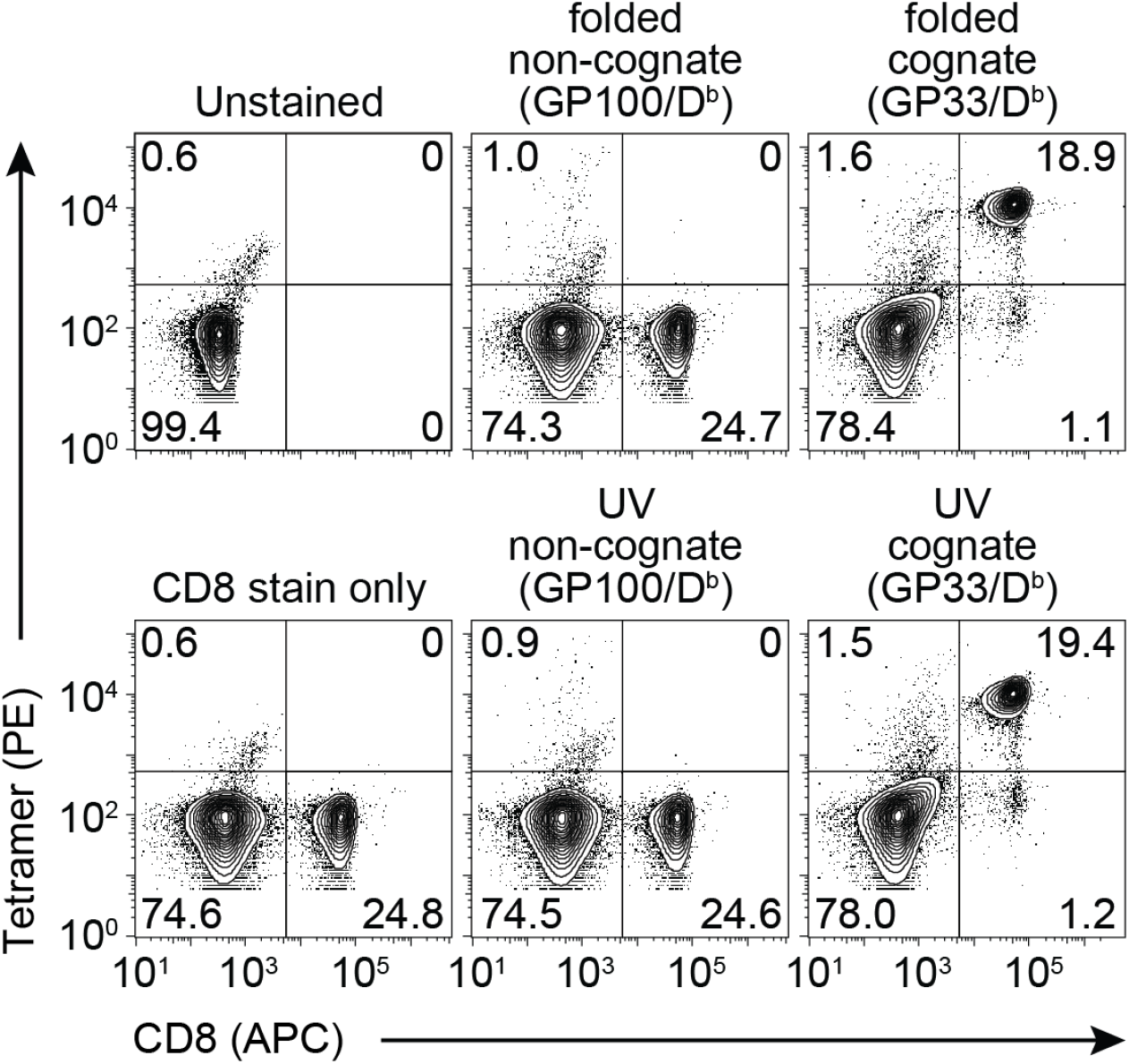
UV-exchanged tetramers bound to P14 CD8 splenocytes comparably to the folded counterpart. Dot plots of flow cytometry analysis for P14 splenocytes stained with cognate (GP33/D^b^) tetramer and non-cognate (GP100/D^b^) tetramer prepared by conventional folding protocol (folded) or UV-mediated ligand exchange protocol (UV).

**Supplementary figure 4.**
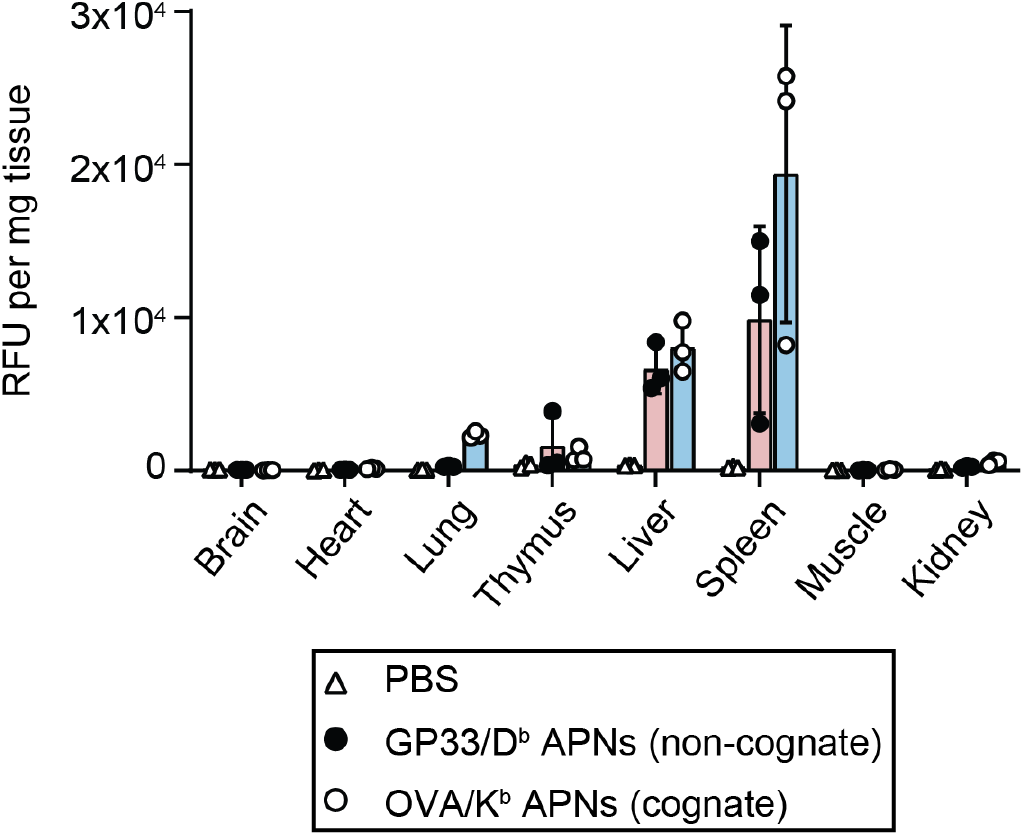
Biodistribution of DiR-labeled APNs. Quantification of DiR fluorescent signal in each organ. Each symbol indicates one measured organ. All data are means ± SD; n = 3 biologically independent mice.

**Supplementary figure 5.**
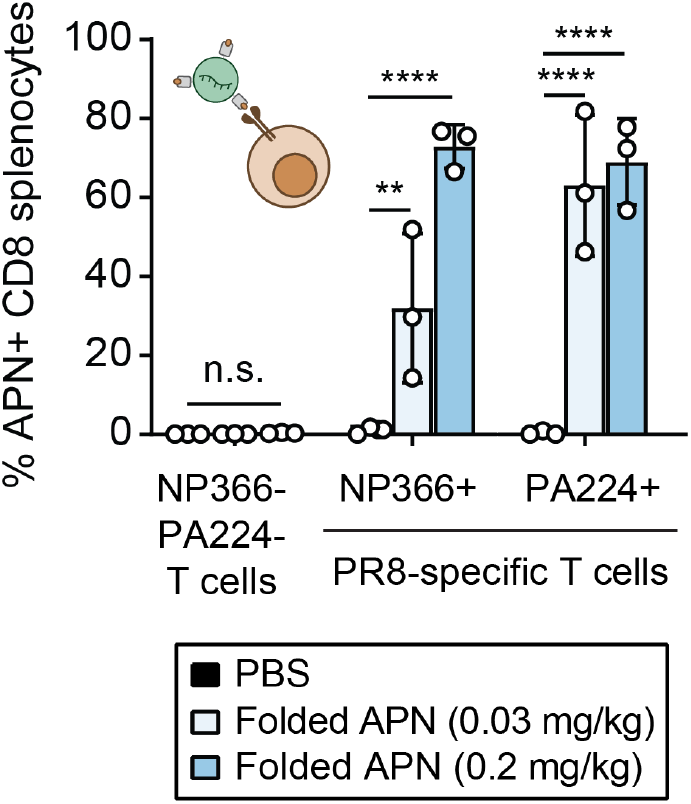
Dose-dependent APN targeting to two immunodominant flu-specific T cells in PR8 model. Infected mice were intravenously injected with a mixture of folded NP366/D^b^ APNs and PA224/D^b^ APNs. n.s. = not significant where *P*>0.9999 between PBS and folded APNs (0.03 mg/kg total mRNA dose) as well as between PBS and folded APNs (0.2 mg/kg) in NP366-PA224-T cells; * * *P* = 0.0033 between PBS and folded APNs (0.03 mg/kg) in NP366+ flu-specific T cell population (NP366+); * * * * *P* < 0.0001; two-way ANOVA and Sidak post-test and correction. All data are means ± SD; n = 3 biologically independent mice.

## REFERENCE

1. Y. Zheng et al., In vivo targeting of adoptively transferred T-cells with antibody- and cytokine-conjugated liposomes. Journal of controlled release 172, 426–435 (2013).

2. D. Schmid et al., T cell-targeting nanoparticles focus delivery of immunotherapy to improve antitumor immunity. Nature communications 8, 1747 (2017).

3. Y. Zheng, L. Tang, L. Mabardi, S. Kumari, D. J. Irvine, Enhancing adoptive cell therapy of cancer through targeted delivery of small-molecule immunomodulators to internalizing or noninternalizing receptors. ACS Nano 11, 3089–3100 (2017).

4. Y.-S. S. Yang et al., Targeting small molecule drugs to T cells with antibody-directed cell-penetrating gold nanoparticles. Biomater Sci 7, 113–124 (2019).

5. S. Ramishetti et al., Systemic gene silencing in primary t lymphocytes using targeted lipid nanoparticles. ACS Nano 9, 6706–6716 (2015).

6. T. T. Smith et al., In situ programming of leukaemia-specific T cells using synthetic DNA nanocarriers. Nature Nanotechnology 12, 813 (2017).

7. R. Kedmi et al., A modular platform for targeted RNAi therapeutics. Nature Nanotechnology 13, 214–219 (2018).

8. N. N. Parayath, S. B. Stephan, A. L. Koehne, P. S. Nelson, M. T. Stephan, In vitro-transcribed antigen receptor mRNA nanocarriers for transient expression in circulating T cells in vivo. Nature communications 11, 6080 (2020).

9. F.-Y. Su, Q. D. Mac, A. Sivakumar, G. A. Kwong, Interfacing biomaterials with synthetic T cell immunity. Advanced Healthcare Materials 10, 2100157 (2021).

10. S. N. Quayle et al., CUE-101, a Novel E7-pHLA-IL2-Fc Fusion Protein, Enhances tumor antigen-specific T-Cell activation for the treatment of hpv16-driven malignancies. Clinical Cancer Research 26, 1953–1964 (2020).

11. A. W. Woodham et al., In vivo detection of antigen-specific CD8+ T cells by immuno-positron emission tomography. Nature Methods 17, 1025–1032 (2020).

12. D. G. Millar et al., Antibody-mediated delivery of viral epitopes to tumors harnesses CMV-specific T cells for cancer therapy. Nature Biotechnology, (2020).

13. J. P. Sefrin et al., Sensitization of Tumors for Attack by Virus-Specific CD8+ T-Cells Through Antibody-Mediated Delivery of Immunogenic T-Cell Epitopes. Front Immunol 10, (2019).

14. X. Clemente-Casares et al., Expanding antigen-specific regulatory networks to treat autoimmunity. Nature 530, 434–440 (2016).

15. S. Singha et al., Peptide–MHC-based nanomedicines for autoimmunity function as T-cell receptor microclustering devices. Nature Nanotechnology 12, 701–710 (2017).

16. A. Tarke et al., Comprehensive analysis of T cell immunodominance and immunoprevalence of SARS-CoV-2 epitopes in COVID-19 cases. Cell Rep Med 2, 100204 (2021).

17. M. Toebes et al., Design and use of conditional MHC class I ligands. Nature medicine 12, 246–251 (2006).

18. M. H. Gee et al., Antigen identification for orphan T Cell receptors expressed on tumor-infiltrating lymphocytes. Cell 172, 549-563.e516 (2018).

19. B. Rodenko et al., Generation of peptide-MHC class I complexes through UV-mediated ligand exchange. Nature protocols 1, 1120–1132 (2006).

20. J. J. Luimstra et al., A flexible MHC class I multimer loading system for large-scale detection of antigen-specific T cells. The Journal of experimental medicine 215, 1493–1504 (2018).

21. S. K. Saini et al., Dipeptides catalyze rapid peptide exchange on MHC class I molecules. Proceedings of the National Academy of Sciences 112, 202–207 (2015).

22. S. A. Overall et al., High throughput pMHC-I tetramer library production using chaperone-mediated peptide exchange. Nature communications 11, 1909 (2020).

23. A. K. Bentzen et al., Large-scale detection of antigen-specific T cells using peptide-MHC-I multimers labeled with DNA barcodes. Nature Biotechnology 34, 1037–1045 (2016).

24. Multi-discipline review: patisiran. Population PK and/or PD analyses (2018 https://www.accessdata.fda.gov/drugsatfda_docs/nda/2018/210922Orig1s000MultiR.pdf.

25. P. M. Tiwari et al., Engineered mRNA-expressed antibodies prevent respiratory syncytial virus infection. Nature communications 9, 3999 (2018).

26. K. Sou, T. Endo, S. Takeoka, E. Tsuchida, Poly(ethylene glycol)-modification of the phospholipid vesicles by using the spontaneous incorporation of poly(ethylene glycol)-lipid into the vesicles. Bioconjugate Chemistry 11, 372–379 (2000).

27. Y. Wang, L. Miao, A. Satterlee, L. Huang, Delivery of oligonucleotides with lipid nanoparticles. Advanced drug delivery reviews 87, 68–80 (2015).

28. P. Agarwal, C. R. Bertozzi, Site-specific antibody–drug conjugates: the nexus of bioorthogonal chemistry, protein engineering, and drug development. Bioconjugate Chemistry 26, 176–192 (2015).

29. M. Shen, J. Rusling, C. K. Dixit, Site-selective orientated immobilization of antibodies and conjugates for immunodiagnostics development. Methods 116, 95–111 (2017).

30. M. T. Puglielli et al., In vivo selection of a lymphocytic choriomeningitis virus variant that affects recognition of the GP33-43 epitope by H-2Db but not H-2Kb. Journal of virology 75, 5099–5107 (2001).

31. J. A. Whelan et al., Specificity of CTL interactions with peptide-MHC class I tetrameric complexes is temperature dependent. Journal of immunology 163, 4342–4348 (1999).

32. J. D. Altman et al., Phenotypic analysis of antigen-specific T lymphocytes. Science 274, 94–96 (1996).

33. G. M. Grotenbreg et al., Discovery of CD8+ T cell epitopes in Chlamydia trachomatis infection through use of caged class I MHC tetramers. Proceedings of the National Academy of Sciences of the United States of America 105, 3831–3836 (2008).

34. S. N. Mueller et al., Qualitatively different memory CD8+ T cells are generated after lymphocytic choriomeningitis virus and influenza virus infections. Journal of immunology (Baltimore, Md. : 1950) 185, 2182–2190 (2010).

35. K. E. Pauken et al., The PD-1 Pathway Regulates Development and Function of Memory CD8 T Cells following Respiratory Viral Infection. Cell reports 31, 107827 (2020).

36. T. Ichinohe et al., Microbiota regulates immune defense against respiratory tract influenza A virus infection. Proc Natl Acad Sci U S A 108, 5354–5359 (2011).

37. B. J. Laidlaw et al., Cooperativity between CD8+ T cells, non-neutralizing antibodies, and alveolar macrophages is important for heterosubtypic influenza virus immunity. PLoS pathogens 9, e1003207–e1003207 (2013).

38. S. N. Mueller, W. A. Langley, E. Carnero, A. García-Sastre, R. Ahmed, Immunization with live attenuated influenza viruses that express altered NS1 proteins results in potent and protective memory CD8+ T-cell responses. Journal of virology 84, 1847–1855 (2010).

39. T. Wu et al., Quantification of epitope abundance reveals the effect of direct and cross-presentation on influenza CTL responses. Nature communications 10, 2846 (2019).

40. D. Rosenblum, N. Joshi, W. Tao, J. M. Karp, D. Peer, Progress and challenges towards targeted delivery of cancer therapeutics. Nature communications 9, 1410 (2018).

41. P. M. Cevaal et al., In Vivo T Cell-Targeting Nanoparticle Drug Delivery Systems: Considerations for Rational Design. ACS Nano 15, 3736–3753 (2021).

42. J. Nikolich-Zugich, M. K. Slifka, I. Messaoudi, The many important facets of T-cell repertoire diversity. Nat Rev Immunol 4, 123–132 (2004).

43. P. J. de Vos van Steenwijk et al., An Unexpectedly Large Polyclonal Repertoire of HPV-Specific T Cells Is Poised for Action in Patients with Cervical Cancer. Cancer research 70, 2707–2717 (2010).

44. R. S. Andersen et al., Dissection of T-cell antigen specificity in human melanoma. Cancer research 72, 1642–1650 (2012).

45. The U.S. National Institute of Allergy and Infectious Diseases, National Institutes of Health. (2021), Immune Epitope Database and Analysis Resource.

46. M. D. Catalina, J. L. Sullivan, K. R. Bak, K. Luzuriaga, Differential evolution and stability of epitope-specific CD8(+) T cell responses in EBV infection. Journal of immunology 167, 4450–4457 (2001).

47. R. Elkington et al., Ex vivo profiling of CD8+-T-cell responses to human cytomegalovirus reveals broad and multispecific reactivities in healthy virus carriers. Journal of virology 77, 5226–5240 (2003).

48. A. H. Bakker et al., Conditional MHC class I ligands and peptide exchange technology for the human MHC gene products HLA-A1, -A3, -A11, and -B7. Proc Natl Acad Sci U S A 105, 3825–3830 (2008).

49. P. A. Chandran et al., A simple and rapid method for quality control of major histocompatibility complex-peptide monomers by flow cytometry. Front Immunol 8, 96 (2017).

50. C. S. Eberhardt et al., Functional HPV-specific PD-1+ stem-like CD8 T cells in head and neck cancer. Nature 597, 279–284 (2021).

51. W. Zhao, X. Sher, Systematically benchmarking peptide-MHC binding predictors: From synthetic to naturally processed epitopes. PLoS Comput Biol 14, e1006457 (2018).

52. K. Sugata et al., Affinity-matured HLA class II dimers for robust staining of antigen-specific CD4+ T cells. Nature Biotechnology 39, 958–967 (2021).

53. N. Veiga et al., Cell specific delivery of modified mRNA expressing therapeutic proteins to leukocytes. Nature communications 9, 4493 (2018).

54. T. Ishida, D. L. Iden, T. M. Allen, A combinatorial approach to producing sterically stabilized (Stealth) immunoliposomal drugs. FEBS Letters 460, 129–133 (1999).

55. A.L. Lainé et al., Conventional versus stealth lipid nanoparticles: Formulation and in vivo fate prediction through FRET monitoring. Journal of Controlled Release 188, 1–8 (2014).

56. S. N. Dahotre, A. M. Romanov, F.-Y. Su, G. A. Kwong, Synthetic antigen-presenting cells for adoptive t cell therapy. Advanced Therapeutics 4, 2100034 (2021).

57. S. N. Dahotre, Y. M. Chang, A. M. Romanov, G. A. Kwong, DNA-Barcoded pMHC tetramers for detection of single antigen-specific T Cells by digital PCR. Analytical Chemistry 91, 2695–2700 (2019).

58. S. N. Dahotre, Y. M. Chang, A. Wieland, S. R. Stammen, G. A. Kwong, Individually addressable and dynamic DNA gates for multiplexed cell sorting. Proceedings of the National Academy of Sciences 115, 4357–4362 (2018).

59. G. A. Kwong et al., Modular Nucleic Acid Assembled p/MHC Microarrays for Multiplexed Sorting of Antigen-Specific T Cells. Journal of the American Chemical Society 131, 9695–9703 (2009).

